# A complete chromosome substitution mapping panel reveals genome-wide epistasis in Arabidopsis

**DOI:** 10.1101/436154

**Authors:** Cris L. Wijnen, Ramon Botet, José van de Belt, Laurens Deurhof, Hans de Jong, C. Bastiaan de Snoo, Rob Dirks, Martin P. Boer, Fred A. van Eeuwijk, Erik Wijnker, Joost J.B. Keurentjes

## Abstract

Chromosome substitution lines (CSLs) are tentatively supreme resources to investigate non-allelic genetic interactions. However, the difficulty of generating such lines in most species largely yielded imperfect CSL panels, prohibiting a systematic dissection of epistasis. Here, we present the development and use of a unique and complete panel of CSLs in *Arabidopsis thaliana*, allowing the full factorial analysis of epistatic interactions. A first comparison of reciprocal single chromosome substitutions revealed a dependency of QTL detection on different genetic backgrounds. The subsequent analysis of the complete panel of CSLs enabled the mapping of the genetic interactors and identified multiple two- and three-way interactions for different traits. Some of the detected epistatic effects were as large as any observed main effect, illustrating the impact of epistasis on quantitative trait variation. We, therefore, have demonstrated the high power of detection and mapping of genome-wide epistasis, confirming the assumed supremacy of comprehensive CSL sets.

**One sentence summary:** Development of a complete panel of chromosome substitution lines enables high power mapping of epistatic interactions in *Arabidopsis thaliana*.

## Main Text

The identification of genetic factors involved in the regulation of quantitative traits is conventionally performed by linkage analysis of genotype-phenotype relationships in segregating mapping populations (*1, 2*). Traditional mapping populations are typically the result of random recombination and segregation of two genotypes in the offspring of an intraspecific cross. Such an approach, however, suffers from a number of inherent complicating factors. These include, amongst others, the simultaneous segregation of multiple quantitative trait loci (QTL) and genetic interactions between them, features that are characteristic for complex polygenic traits. As a result, conventional mapping populations, such as recombinant inbred lines (RILs), require a large collection of segregating lines to obtain sufficient statistical power to unequivocally detect QTLs and epistasis (*1, 3*). Alternatively, chromosome substitution lines may offer a powerful mapping resource for the systematic dissection of epistatic interactions (*4, 5*).

Chromosome substitution lines (CSLs), a.k.a. consomic strains in non-plant species, differ from established mapping populations by their lack of intra-chromosomal recombination. Consequently, CSLs consist of an assembly of non-recombinant chromosomes, each derived from either one of two genetically different parents (*5, 6*). Genetic variation in CSL populations thus depends exclusively on the reshuffling of complete genotypically distinct chromosomes. As a consequence, the maximum size of chromosome substitution panels, *i.e.* all possible combinations of chromosomes, is finite (2n, where n is the haploid chromosome number), depending solely on the chromosome number of the subjected species. Complete sets of CSLs offer the advantage of fully balanced allele frequency distributions, providing equal haplotype class sizes in epistatic analyses, and a relatively small population size for species with low chromosome numbers, allowing high line replication in experiments. To date, a nearly complete set of CSLs has only been established in *Drosophila melanogaster*, due to the ease of generating CSLs and the limited chromosome number in this species (*7*). However, for most other species, complete sets of CSLs are notoriously difficult to generate using conventional backcross approaches and, despite their promises, only a very limited number of CSLs in just a handful of vertebrate and plant species have been developed (*6–10*). Moreover, all these existing panels consist of CSLs with an introgression of only a single donor chromosome in a recurrent genetic background, which considerably restricts the analysis of epistatic interactions. Nonetheless, single chromosome substitution lines (sCSLs) allow the straightforward detection of additive main effects of introgressed chromosomes, while a deviation of the cumulative sum of these effects from the wild type donor phenotype might indicate the presence of epistatic interactions (*4*). However, the exact strength and genetic architecture of epistasis can only be decomposed by investigating the combined effect of multiple chromosome substitutions.

The recently emerged reverse breeding technology in Arabidopsis determined a major step forward for the development of CSLs (*2, 11*). This approach makes use of the random segregation of non-recombinant chromosomes to the gametes of achiasmatic hybrids, resulting from the transgenic repression of recombination. These gametes are then converted into haploid offspring through crossing to a haploid inducer line (*12*). Finally, the haploid progeny, which consist of an assembly of non-recombinant chromosomes, each derived from either one of the two parents of the initial hybrid, is converted into immortal doubled haploids (DHs). DH seeds occur spontaneously in haploid plants at a low frequency either due to the merging of incidental unreduced gametes that arise by chance, or by somatic doubling. The CSLs produced in this way are now normal diploids containing completely homozygous pairs of chromosomes descending from either parent but in a cytoplasmic background of the haploid inducer line. In Arabidopsis, encompassing five chromosome pairs, a complete biparental panel of all possible CSLs comprises 32 (=2^5^) different genotypes (Fig. 1).

**Fig. 1:**
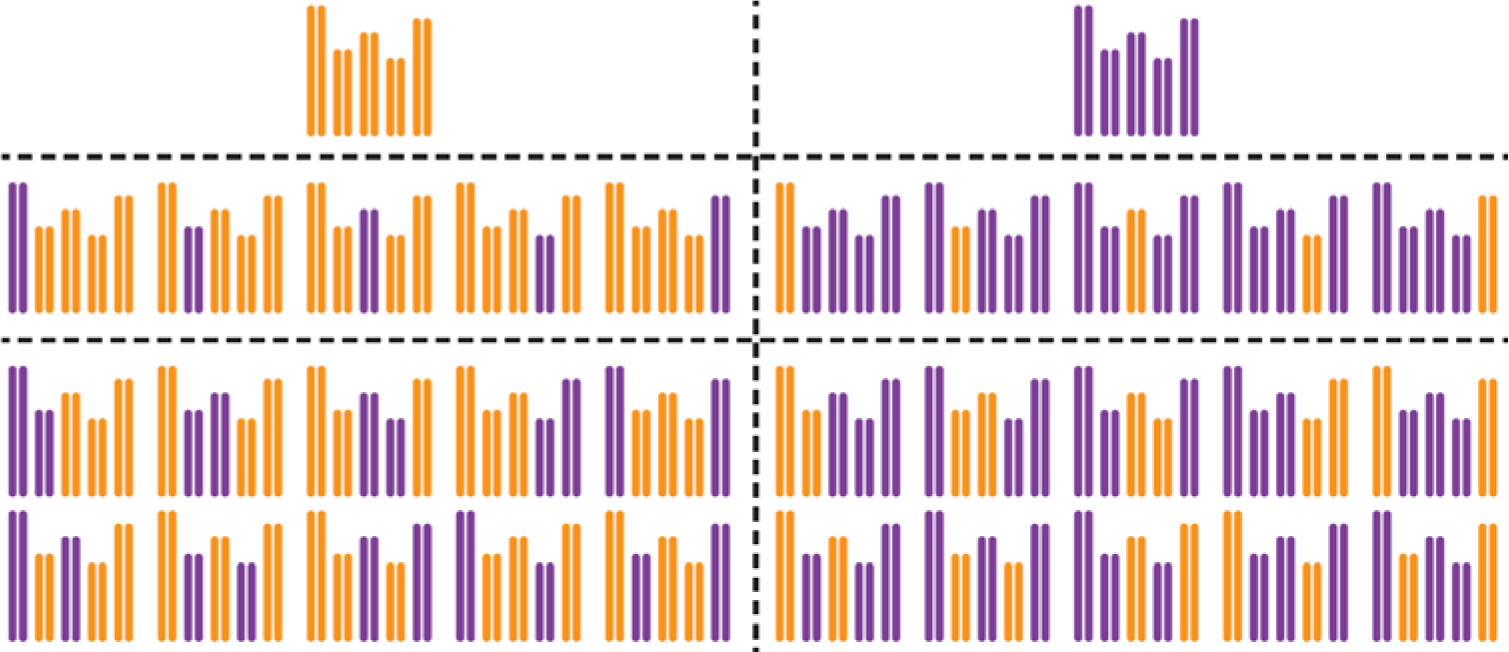
A complete set of Arabidopsis *thaliana* chromosome substitution lines. The complete panel of 32 CSLs can be divided into two reciprocal recurrent backgrounds (vertical dashed line) and subgroups of parental genotypes, single CSLs and CSLs in which two chromosomes are exchanged (horizontal dashed lines). Arabidopsis genomes of each of the CSLs are represented by five homozygous chromosomes derived from either the Col-0 (orange) or L*er* (purple) accession.

Here, we report on the construction and application of such a complete set of CSLs resulting from a cross between the Arabidopsis accessions Columbia-0 (Col-0) and Landsberg *erecta* (L*er*) (Fig. 1 and Fig. S1). Two of the 32 CSLs resemble the identical genotype of the original parents, albeit both in the cytoplasmic background of the inducer line now (*viz*. Col-0). However, ten (2×5, reciprocally) CSLs contain a single substituted chromosome (sCSL), whereas the other twenty (2×10, reciprocally) CSLs contain multiple substituted chromosomes. To demonstrate the enhanced power of complete CSL panels in genetic mapping and epistatic analyses, the complete panel was grown in a climate-controlled growth chamber under short day conditions. In order to compare the performance of CSL mapping with conventional linkage analysis a population of RILs derived from the same accessions was grown simultaneously (*13*). All plants were phenotyped for flowering time (days after germination) and main stem length (mm) at the moment of opening of the first flower.

In accordance with the use of conventional consomic strains the additive effect of a single substituted chromosome in comparison to the non-substituted recurrent parental genotype can be analysed. Moreover, since we have generated sCSLs in both recurrent parental backgrounds we can also specifically assess the contribution of epistatic effects to phenotypic variation (*14*). Using a regression model obtained via a backward elimination procedure, significant effects on flowering time were detected for the substitution of the L*er* chromosomes 2, 3, 4 and 5 in the Col background (Fig. 2A; Table 1; Table S1). Similarly, significant effects on main stem length were observed for the substitution of the L*er* chromosomes 1, 2, 3 and 5 in the Col background (Fig. 2B; Table 1; Table S1). However, in contrast to the reciprocal exchange, the substitution of Col chromosome 3 in a L*er* background displayed no significant effect on flowering time, while the substitution of chromosome 1 did. Likewise, the substitution of the L*er* chromosomes 1 and 3 in a Col background had no significant effect on main stem length, while substitution of these chromosomes in a L*er* background led to significant differences. In addition to these qualitative background differences, the quantitative effect sizes of the reciprocal substitutions differed substantially. Although the largest effect on main stem length was caused by a substitution of chromosome 2 in both backgrounds, the size of the effect differed approximately four-fold. Furthermore, flowering time was mainly affected by substitution of chromosome 5 in the Col-0 background, whereas the largest effect in the L*er* background was obtained by the substitution of chromosome 2. These differences indicate both qualitative as well as quantitative interaction effects of single chromosome substitutions with the remainder of the genome. Indeed, when the substituted chromosomes and the recurrent background were both included in the regression model, significant interactions of most chromosomes with their background were detected for both traits (Fig. 2A-B; Table 1).

Strikingly, the number of QTLs detected in the RIL population using conventional composite interval mapping (CIM) was much lower than in the sCSL panel (Table S1), as was also previously observed for rodents (*15*). For flowering time two significant QTLs were detected on chromosome 2 and an additional one on chromosome 1 but no QTLs were detected on any of the other three chromosomes, consistent with previous studies (*16*) (Fig. 2C). Furthermore, variation in main stem length in the RIL population is largely explained by a single QTL on chromosome 2, most likely reflecting allelic variation of the *ERECTA* locus (*17*) (Fig. 2D). So, in concordance with studies of sCSLs in rodents, QTL detection in our CSL population outperformed traditional linkage mapping in RILs for all traits analysed.

**Table 1:** Regression models for different CSL populations explaining variation in flowering time and main stem length. Populations consist of CSLs with only a single substituted chromosome in a particular background plus their recurrent parent, a set of all sCSLs plus recurrent parents, or the complete set of CSLs, including parental genotypes. Regression models contain only backward selected parameters significantly contributing to explained variance. The parameters *Chr1*, *Chr2*, *Chr3*, *Chr4* and *Chr5* denote additive effects of individual chromosomes whereas *BG* denotes background effects. Parameter components separated by a colon indicate interaction effects.

| Population | Background | Flowering time | Main stem length |
| --- | --- | --- | --- |
| 5 sCSLs +<br>Col parent | Col | *Chr2 + Chr3 + Chr4 +*<br>*Chr5* | *Chr1 + Chr2 + Chr3 +*<br>*Chr5* |
| 5 sCSLs +<br>Ler parent | Ler | *Chr1 + Chr2 + Chr4 +*<br>*Chr5* | *Chr2 + Chr5* |
| 10 sCSLs +<br>both parents | Col + Ler | *Chr1 + Chr2 + Chr3 +*<br>*Chr4 + Chr5 + BG +*<br>*Chr3:BG + Chr4:BG +*<br>*Chr5:BG* | *Chr1 + Chr2 + Chr3 +*<br>*Chr5 + BG + Chr2:BG*<br>*+ Chr3:BG* |
| 32 CSLs | Col + Ler | *Chr1 + Chr2 + Chr3 +*<br>*Chr5 + Chr1:Chr3 +*<br>*Chr1:Chr5 +*<br>*Chr3:Chr5* | *Chr1 + Chr2 + Chr3 +*<br>*Chr5 + Chr1:Chr2 +*<br>*Chr1:Chr5 + Chr2:Chr5*<br>*+ Chr3:Chr5 +*<br>*Chr1:Chr2:Chr5* |

**Fig. 2:**
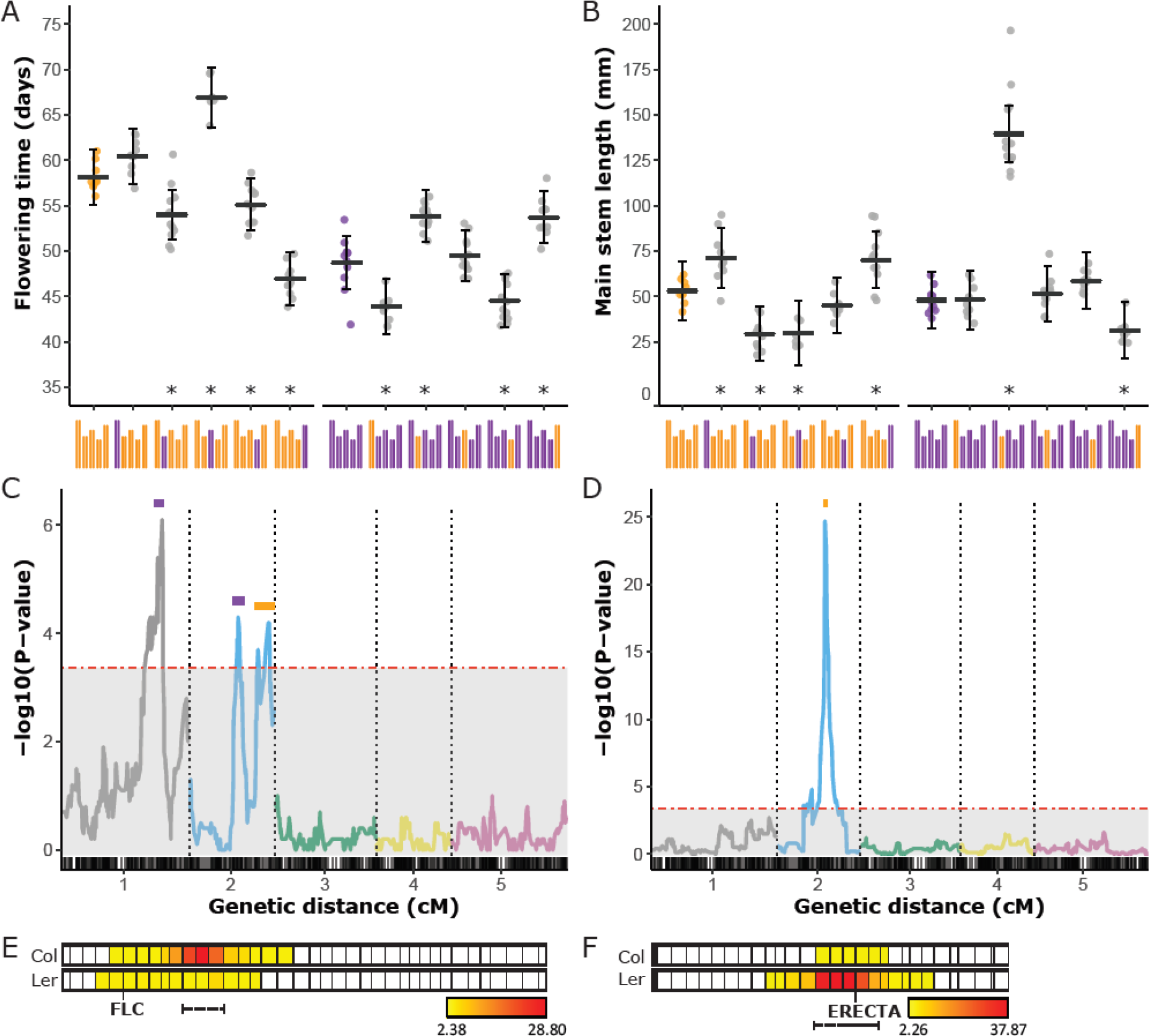
Mapping and validation of single chromosome substitution effects. A-B) Flowering time (A) and main stem length (B) of sCSLs and their recurrent parents. Each dot represents the spatial corrected trait value of an individual of the genotype indicated below the x-axis. Horizontal bars indicate BLUPs with 95% confidence intervals shown as vertical bars (Table S15). Asterisks denote significant effects. C-D) QTL plots for flowering time (C) and main stem length (D) as mapped in a RIL population.–log10(P) values for each chromosome are displayed in different colours, while the horizontal red dashed line represents the significance threshold. Support intervals for the QTLs are indicated by coloured bars according to effect sign (orange: +Col-0, and purple: +L*er*). The x-axis indicates chromosome numbers below a rug profile of the marker positions in cM distance. E-F) Heatmap plots of the effect strength of reciprocal chromosome five introgression NILs on flowering time (E) and chromosome two introgression NILs on main stem length (F). In both panels the upper row represents NIL mapping in a Col background, whereas the lower row represents NIL mapping in a *Ler* background. Vertical lines indicate marker positions in cM. Color intensity from yellow to red specifies the strength of significant effects. Dashed lines below the heatmap indicate support intervals. FLC and ERECTA indicate the position of obvious candidate genes explaining variation in flowering time and main stem length, respectively.

Despite the high detection power, CSLs inherently offer a low resolution since QTLs can only be mapped to entire chromosomes due to the lack of recombination. To overcome this drawback a reciprocal genome-wide coverage set of near-isogenic lines (NILs) was generated. These were produced by backcrossing sCSLs to one of the recurrent parental accessions and subsequent DH production of recombinant F1 gametes, as described for the generation of CSLs. In total 413 NILs with either a single or multiple introgressions were generated of which 219 contained a L*er* introgression in a Col background and 194 contained a Col introgression in a L*er* background, as determined by marker-assisted genotyping (Table S3). This genetic resource serves to validate and fine-map detected QTLs in the CSLs and confirm possible epistatic interactions with the genetic background.

To demonstrate the complementing value of this NIL population, a subset of reciprocal NILs covering the chromosomes 2 and 5 were grown in similar conditions as the CSLs and RILs. The substitution of chromosome 2 had the largest effect on main stem length, with two-fold longer stems in genotypes carrying a Col chromosome 2 (Fig. 2E). Fine-mapping of this chromosome in the reciprocal NILs resulted in a support interval of 9.1-16.5 Mbp for the Col set, while this was much narrower in the L*er* set, 9.9-11.3 Mbp. This coincides well with the support interval of the QTL mapped in the RIL population (11.1-11.7 Mbp, Fig. 2D) and covers the position of the obvious candidate gene *ERECTA* at 11.2 Mbp. A similar resolution, support interval 7.3-8.8 Mbp and 8.0–9.7 Mbp for Col and L*er* NILs respectively, could be obtained for the fine-mapping of the chromosome 5 QTL for flowering time. No obvious candidate genes are positioned within this support interval although the strong *FLOWERING LOCUS C* (*FLC*) was located on the same chromosome arm (Fig. 2F). Surprisingly, despite a ten-day delay in flowering time in sCSLs in which a L*er* chromosome 5 is substituted in a Col background, this QTL was not detected in the RILs (Fig. 2C).

An interesting observation from the analysis of the reciprocal NIL sets is the difference in mapping power. The effect on flowering time of a L*er* chromosome 5 substitution in a Col background (ΔFT=−7.4 days) is much larger than *vice versa* (ΔFT=+4.5 days). Likewise, the effect on main stem length of a Col chromosome 2 substitution in a L*er* background is almost eightfold larger than *vice versa*. These differences might reflect discrepancies in effect sizes relative to the recurrent parent’s trait value, which might be the result of an accumulation of additive effects, or could indicate a dependency on epistatic interactions. Although the limited set of reciprocal sCSLs also indicates the presence of epistasis, both chromosome 2 and 5 were identified to interact with the background in determining main stem length and flowering time, respectively, the specific origin of these genetic interactions can only be identified by comparing CSLs with multiple substituted chromosomes.

The importance of genetic interactions, relative to the additive effects of single loci, on the phenotypic expression of a trait is part of a long lasting debate (*18–20*) and multiple studies have reported on models including epistasis that explain more variation (*21*) and have a better predictive power (*22*) compared to models including only main effects. However, the unbiased testing of epistasis as a source of natural variation is statistically challenging since increasing levels of interaction decrease the number of observations for each genotypic class, which drains the power to detect interacting loci. Furthermore, in most standard mapping populations undetected QTLs are added to the error term. Finally, overfitting of a model can become a problem due to the close to an infinite number of allelic combinations in a segregating recombinant biparental population. Therefore, most statistical models only include main additive effects and the interactions between them, leaving part of the heritable variation unexplained (*18*). Completely balanced CSL panels, however, offer the unique opportunity to analyse the relatively limited number of all possible genotypic combinations in a full factorial design and as such provide a more realistic view on the complexity of quantitative trait regulation.

Since clear indications of genetic interactions between chromosomes were obtained from the analysis of reciprocal CSLs and NILs, a regression analysis using a backward elimination strategy on data of the complete CSL panel (Fig. 3A-B) was performed to quantify the contribution of epistasis to the phenotype. Using a similar regression approach as was used to test the sCSLs for background interactions, significant chromosome interactions were included in the final model. For flowering time, significant two-way interactions were detected between chromosome 1 and 3, 1 and 5, and 3 and 5 (Fig. 3C-E), which partly explain the major effect of genotypic variation of chromosome 5 (Fig. 3A). For main stem length a significant three-way interaction between the chromosomes 1, 2 and 5 was detected, while a significant two-way interaction was detected between chromosome 3 and 5 (Fig. 3F-G).

**Fig. 3:**
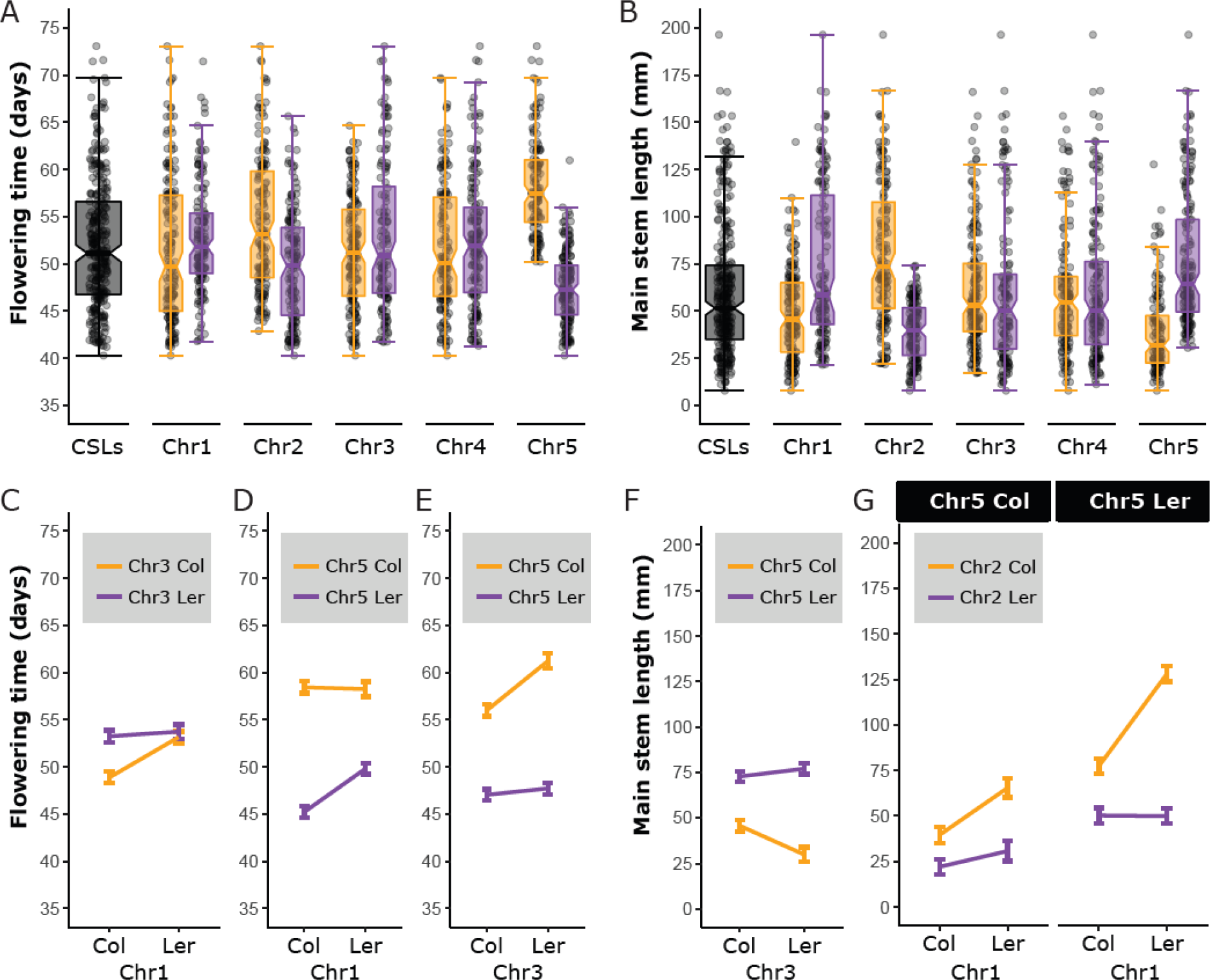
Detection of interchromosomal interaction effects in a complete CSL panel. A-B) Notched box-and-whisker plots of the complete panel of CSLs for flowering time (A) and main stem length (B). Each dot represents the spatial corrected trait value of an individual plotted in relation to all other individuals (grey boxes) or categorized according to its genotype for the chromosome indicated on the X-axis (orange boxes: Col; purple boxes: L*er*). C-G Regression predicted effect plots of epistatic interactions identified with backward selection models. C-E) Two-way interactions explaining variation in flowering time. F) Two-way interaction explaining variation in main stem length. G) Three-way interaction explaining variation in main stem length. Error bars represent the 95% confidence intervals of the predicted effect.

Although in general main effect sizes are considered to be larger than interaction effects, here the interaction effect of three chromosomes on main stem length is of similar size as the most effective substitution of a single chromosome (Fig. 2B). Most notable for this three-way interaction is a more than 65% increase in main stem length of one genotypic class (Chr1^Ler^/Chr2^Col^/Chr5^Ler^) over any of the other seven genotypic classes (Fig. 3G). The importance of epistasis is also demonstrated by a comparison of regression models which either include or exclude epistatic interactions. An inclusive model displays a superior predictive power (R^2^ =; 0.835) over a model in which epistatic interactions are not considered (R^2^ = 0.760; Fig. S2). Finally, the impact that genetic interactions can have on the phenotype is illustrated by a case of antagonistic epistasis between chromosome 3 and 5, where the substitution of a Col chromosome 3 with that of L*er* resulted in opposite effects on main stem length, depending on the genotype of chromosome 5.

Our results show that a relatively large part of the observed variation in the analysed quantitative traits can be explained by epistatic interactions. The power to detect these interactions and estimate their effect sizes is greatly enhanced by analysing a complete panel of CSLs, which also includes lines in which multiple chromosomes are substituted. The notion that even for traits dominated by major effect loci (*e.g. ERECTA* in main stem length) epistatic interactions can be revealed, and given the small size of this population, CSL mapping holds great promises for many other quantitative traits in Arabidopsis. There is no reason to assume that similar results cannot be obtained in other species, although larger genome sizes (*i.e.* higher chromosome numbers) might require the simultaneous substitution of two or more chromosomes.

## Acknowledgements

We like to express our gratitude to F. Becker, G. Stunnenberg, T. Stoker and R. van Genderen of Wageningen University for technical assistance during experimental work.

## Author contributions

R.D., E.W., J.J.B.K. conceptualized experiments; H.d.J., M.P.B., F.A.v.E., E.W., J.J.B.K. acquired funding; M.P.B., F.A.v.E., E.W., J.J.B.K. supervised research activities; C.L.W., R.B., E.W., J.J.B.K. designed experiments; C.L.W., R.B., J.v.d.B., L.D., C.B.d.S. performed experiments; C.L.W., R.B., M.P.B., F.A.v.E. analysed data; C.L.W., R.B., E.W., J.J.B.K. wrote the original draft; and all authors discussed the results and commented on the manuscript.

## Data and materials availability

All data is available in the main text or the supplementary materials. Materials will be donated to the appropriate stock centers.

